# Multi-scale responses of scattering layers to environmental variability in Monterey Bay, California

**DOI:** 10.1101/047902

**Authors:** Samuel S. Urmy, John K. Horne

**Author notes:** Current address: Stony Brook University School of Marine and Atmospheric Sciences, Montauk Hwy, Southampton, NY 11968, USA. Email address (Samuel S. Urmy).

## Abstract

A 38 kHz upward-facing echosounder was deployed on the seafloor at a depth of 875 m in Monterey Bay, CA, USA (36° 42.748’ N, 122° 11.214’ W) from 27 February 2009 to 18 August 2010. This 18-month record of acoustic backscatter was compared to oceanographic time series from a nearby data buoy to investigate the responses of animals in sound-scattering layers to oceanic variability at seasonal and sub-seasonal time scales. Pelagic animals, as measured by acoustic backscatter, moved higher in the water column and decreased in abundance during spring upwelling, attributed to avoidance of a shoaling oxycline and advection offshore. Seasonal changes were most evident in a non-migrating scattering layer near 500 m depth that disappeared in spring and reappeared in summer, building to a seasonal maximum in fall. At sub-seasonal time scales, similar responses were observed after individual upwelling events, though they were much weaker than the seasonal relationship. Correlations of acoustic backscatter with oceanographic variability also differed with depth. Backscatter in the upper water column decreased immediately following upwelling, then increased approximately 20 days later. Similar correlations existed deeper in the water column, but at increasing lags, suggesting that near-surface productivity propagated down the water column at 10-15 m d^‒1^, consistent with sinking speeds of marine snow measured in Monterey Bay. Sub-seasonal variability in backscatter was best correlated with sea-surface height, suggesting that passive physical transport was most important at these time scales.

## 1. Introduction

Physical variability is a fundamental feature of the ocean’s pelagic habitat, and the responses of pelagic organisms to this variability play a large role in determining their distribution, abundance, and survival. As such, understanding the effects of physical change and variability on ocean life has long been recognized as a central challenge in oceanography and marine ecology. Physical variability can influence organisms at the individual or population level, and its effects can be direct (e.g. advection) or indirect (e.g production fertilized by upwelled nutrients).

“Physical-biological coupling” has most often been studied in the plankton, whose abundance and distribution are closely tied to physical variability (Platt and Denman, 1975). However, because ocean currents are variable across a wide range of spatial and temporal scales, the division between plankton and nekton (Haeckel, 1890) is somewhat arbitrary. Coupling to physical processes is thus expected to extend from zooplankton to micronekton: animals roughly 2-10 cm in length with swimming abilities in between those of drifting plankton and freely swimming nekton (Brodeur and Yama-mura, 2005). These include krill, pelagic shrimps, small squids, and fishes such as myctophids.

Assemblages of micronekton are important constituents of deep scattering layers (DSLs, Dietz 1948; Barham 1956). DSLs are layers of elevated animal biomass, and consequently acoustic backscattering, which are found worldwide in the ocean’s mesopelagic zone, ≈ 200 to 1,000 m below the surface. Many undergo diel vertical migration (DVM) of several hundred meters to feed near the surface each night (Dietz, 1948; Hays, 2003). Micronekton, from both the meso‐ and epipelagic (0-200 m depth), are important food resources for a variety of larger fish, birds, and marine mammals and are important carriers of energy, both up the food chain and down the water column. Recent research suggests that the global biomass of small fishes in the DSL is on the order of 10^10^ metric tons, and that they could respire as much as 10% of primary productivity in the deep ocean (Kaartvedt et al., 2012; Irigoien et al., 2014). They are probably also influenced by physical variability, especially in dynamic environments.

The California Current is one such dynamic environment. As in other eastern-boundary systems, Ekman pumping driven by seasonal equatorward-flowing winds brings nutrient-rich water to the surface near the coast, fueling a highly productive ecosystem. Though limited in area, upwelling systems support a disproportionate amount of global fish landings (Pauly and Christensen, 1995), and attract large predators from great distances (Block et al., 2011). The seasonal cycle of upwelling and productivity in the California Current is generally consistent (Pennington and Chavez, 2000), but within any given year upwelling is irregular, supplying nutrients to the food web in episodic pulses. Upwelling is also spatially variable. Mesoscale (10s to 100s of km) squirts, jets, eddies, and coastal waves all introduce variability into the movement of water (Keister and Strub, 2008).

The effects of oceanic variability on micronekton are only now beginning to be understood. Responses of phytoplankton to environmental variability have been well studied at interannual (McGowan et al., 2003), seasonal (Bolin and Abbot, 1963; Service et al., 1998), and sub-seasonal time scales (Service et al., 1998; Legaard and Thomas, 2007). Environmental effects on zooplankton and micronekton have also been studied, though mostly at seasonal and longer time scales (e.g. Roesler and Chelton 1987; McGowan et al. 1996; Brinton and Townsend 2003; Rebstock 2003). Several recent studies have examined spatial relationships between physical features and micronekton distribution. These have regularly found changes in DSL structure and density associated with mesoscale eddies (Kloser et al., 2009; Godø et al., 2012; Fennell and Rose, 2015) and across frontal zones (Opdal et al., 2008; Irigoien et al., 2014; Boersch-Supan et al., 2015). Fewer studies have examined the temporal evolution of DSLs in relation to physical oceanography (Wang et al., 2014), and at sub-seasonal temporal scales, these effects are largely unknown.

This study used a bottom-mounted echosounder in outer Monterey Bay, CA to monitor changes in acoustic backscatter over 18 months. This sampling allowed us to characterize macrozooplankton and micronekton biomass through the entire water column at high temporal and spatial resolutions. We compared these acoustic backscatter measurements to coincident measurements of wind, temperature, and fluorescence as proxies of upwelling, and sea levels, to investigate possible links between sea-surface topography and animal biomass (e.g. Clarke and Dottori 2008). We also examined how these responses varied as a function of depth. We expected a lagged increase in backscatter following upwelling events, due to a combination of animal aggregation, somatic growth, and population increase. A similar response to increased productivity at the surface was expected from animals at depth, but delayed and damped when compared to the response of surface animals. Finally, we estimate the relative importance of different processes in generating the observed physical-biological relationships.

## 2. Methods

### 2.1. Study location

Monterey Bay is a large, open embayment in the central California coast. The Bay’s oceanographic seasons follow those of the California Current, with wind-driven upwelling in spring and early summer, a warm water “oceanic period” in the late summer and fall, and a winter downwelling or “Davidson current” period (Skogsberg et al., 1946; Pennington and Chavez, 2000). Point Año Nuevo, to the north of the Bay, is the source of a persistent upwelling plume that typically trails south across the mouth of the Bay (Rosenfeld et al. 1994, Figure 1). Mean circulation within the bay is counterclockwise, with enhanced productivity in the “upwelling shadows” near shore (Graham et al., 1992).

**Figure 1:**
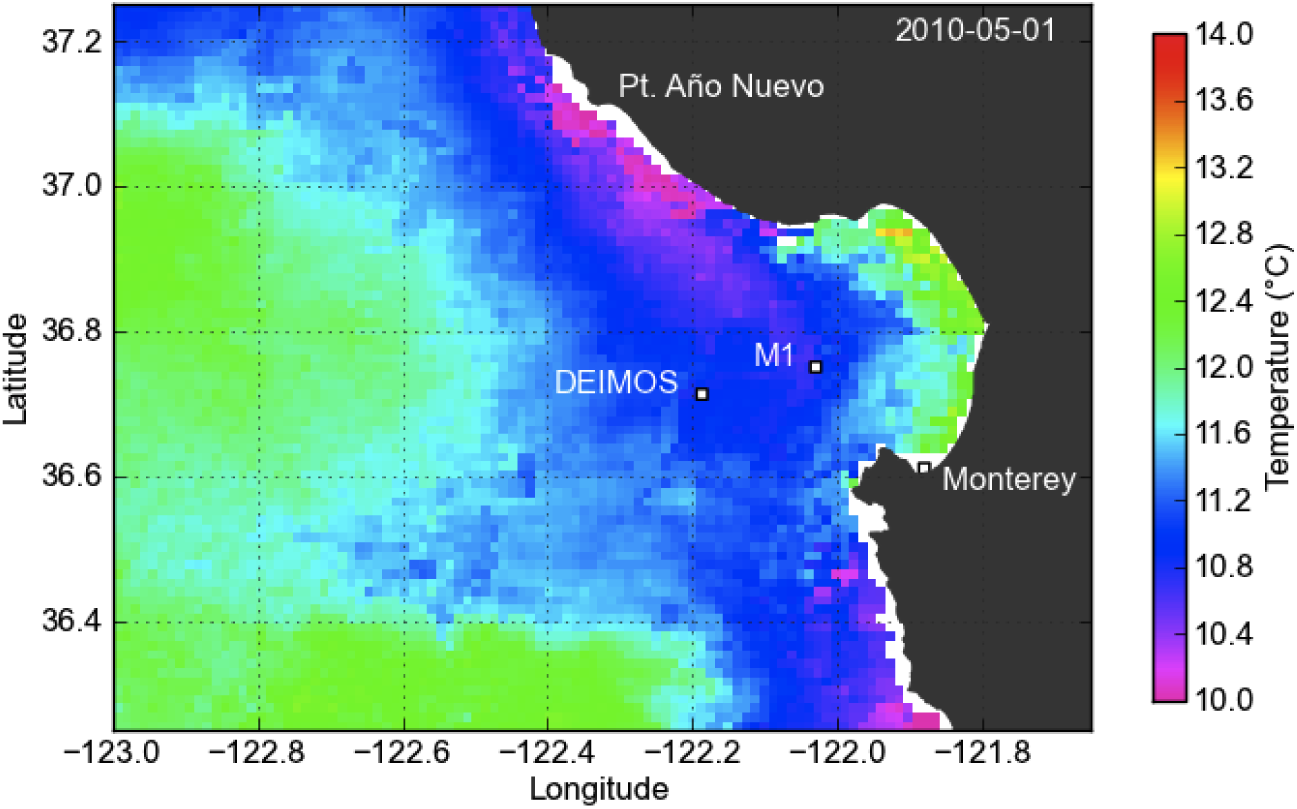
Monterey Bay, showing location of the upward-facing echosounder (DEIMOS) and oceanographic data buoy (M1) used in this study, as well as a typical pattern of sea-surface temperature during the upwelling season (AVHRR 3-day composite, 1 May, 2010).A band of cold, upwelled water is located along the coast, with warmer water offshore and inside the Bay. The coldest waters are near Point Año Nuevo and Point Sur.

### 2.2. Acoustic Data

Animal density through the water column was estimated using a bottom-mounted echosounder. The Deep Echo Integrating Marine Observatory System (DEIMOS) is an acoustic package built around a 38 kHz scientific echosounder (Horne et al., 2010). It was deployed at 875 m depth from February 27, 2009 to August 18, 2010 at the Monterey Accelerated Research System (MARS), a cabled observatory node, located at 36°42.748’ N, 122°11.214’ W on Smooth Ridge, to the north of the Monterey Submarine Canyon. MARS is maintained and operated by the Monterey Bay Aquarium Research Institute (MBARI), and provides continuous power and communications for scientific instruments. Several multi-day gaps in the data were caused by electrical interference, software crashes, and burrowing rodents (Urmy et al., 2012). DEIMOS sampled continuously at 0.2 Hz with a 0.5 m vertical resolution through the water column. DEIMOS was calibrated *in situ* using a standard target (Foote et al., 1987) hung from a float above the transducer during the final 7 weeks of the deployment.

We were not able to take direct samples to identify scattering organisms, so acoustic volume and area backscattering coefficients (*s*_*v*_ and *s*_*a*_, and their logarithmic forms *S*_*v*_ and *S*_*a*_, MacLennan et al. 2002) were used as proxies of animal biomass. This is a reasonable assumption for both single species (Foote, 1983) and mixed communities (Benoit-Bird and Au, 2002).

Acoustic data were processed using Echoview software (version 4.8, Myriax Pty. Ltd. 2010). Background noise was estimated and subtracted using methods described in De Robertis and Higginbottom (2007). A backscatter threshold was applied to eliminate acoustic returns with volume-scattering strengths below ‐90 dB, the approximate backscattering intensity generated by one krill m^‒3^ at 38 kHz (Demer and Conti, 2003). All echograms were visually inspected, and regions with external noise (e.g. ship or ROV noise) were excluded from further analysis. Also excluded were regions within 7 m of the bottom, to eliminate targets in the acoustic near field, and within 10 m of the surface, to avoid integrating bubbles from breaking waves. The mean depth of backscatter was measured using the acoustic center of mass (CM, Urmy et al. 2012).

### 2.3. Oceanographic Data

Time series of wind velocity, sea-surface temperature (SST), and fluorescence, a proxy for chlorophyll (Kirk, 1994), were measured at MBARI’s M1 data buoy, located 15 km ENE of DEIMOS at 36°45’ N, 122° 1.8’ W (Chavez et al. 1997, Figure 1). Service et al. (1998) found that wind velocity at M1 was highly correlated (*R*^2^ = 0.78) with wind velocity at MBARI’s M2 mooring, 18 km WSW of DEIMOS. This correlation held during our study period (*R*^2^ = 0.73), indicating that winds at M1 were representative of those at DEIMOS, located approximately midway between the M1 and M2 moorings. Daily satellite measurements of SST and log chlorophyll-a at DEIMOS and M1, from Level-3 AVHRR and MODIS-Aqua imagery, were also correlated (*R*^2^ = 0.86 and 0.37), giving us confidence that SST and fluorescence measurements at M1 were representative of these values at the DEIMOS site.

Ekman transport of water offshore, an estimate of wind-driven upwelling, was calculated from wind measurements at M1 following Bakun (1973). Offshore Ekman transport was estimated as *M*_*E*_ = *τ*_*a*_/*f*, where *τ*_*a*_ is the alongshore component of the wind stress and *f* is the Coriolis acceleration, equal to 8.326 × 10^‒5^ s^‒1^ at latitude 36°45’ N. Wind stress was calculated as ***τ*** = *ρC*_*d*_ |**u**| **u**, where **u** is wind velocity, *ρ* is the density of air (assumed constant at 1.22 kg m^‒3^), and *C*_*d*_ is a non-dimensional drag coefficient, taken to be 0.0013 (Bakun, 1973; Schwing et al., 1996). Alongshore wind stress was defined as the component parallel to 150°, with positive stresses towards the southeast. Sea-surface height (SSH) was measured by the National Oceanographic and Atmospheric Administration (NOAA) tide gauge in Monterey (http://tidesandcurrents.noaa.gov/geo.shtml?location=monterey). SSH was corrected for the reverse-barometer effect (Chelton and Enfield, 1986) and low-pass filtered with a 25-hour moving average to remove the effects of diurnal and semidiurnal tides.

### 2.4. Analysis

Prior to analysis, values in all time series were averaged into one-day bins. For the acoustic data, these averages used only values within four hours of local noon (i.e. 08:00-16:00) to avoid “blurring” by diel vertical migration. Oceanographic series were averaged over the full 24 hour period.

Correlations between environmental and acoustic variables at the seasonal scale were quantified by calculating Pearson product-moment correlation coefficients between the respective time series. We also quantified the seasonal cycles by fitting composite sinusoids with periods of 12 and 6 months to the data by least squares. This allowed us to estimate the amplitudes and timings of these cycles, even though only 18 months of acoustic data were available. All further analyses used the residuals from these model fits.

We calculated the correlation between oceanographic and acoustic series at time lags from 0 to 90 days—i.e., the cross-correlation function (CCF). We used only the half of the CCF where the oceanographic variable led the acoustic variable, since we were interested in the influence of oceanography on micronekton. This is consistent with “bottom-up” forcing, from physics to primary production and consumption, usually assumed to operate through lower trophic levels (cf. Micheli et al. 1999). Assuming an uncorrelated white-noise null hypothesis, CCFs were considered significant if their absolute value was greater than 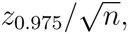, where *z*_0.975_ is the 0.975 quantile of the standard normal distribution and *n* is the number of observations in each time series (Brockwell and Davis, 2002). We calculated CCFs of temperature and fluorescence with wind stress to check the time lag of phytoplankton blooms behind upwelling events. We then calculated the lagged correlation of upwelling wind stress, temperature, fluorescence, and sea level with depth-integrated backscatter (*s*_*a*_) and the CM. We also calculated the lagged correlation of backscatter, at each depth in the water column, with the oceanographic series.

To test the predictive power of the bio-physical relationships, we built a statistical model for backscatter in the top 300 m of the water column, representing prey available to surface-diving and epipelagic predators. We regressed *S*_*a*_, integrated from 0-300 m, on the values of the four environmental variables at their best-correlated lag below 30 days, selecting significant covariates using a backwards-deletion procedure. An autoregressive (AR) model accounted for autocorrelation in the residuals. We computed the corrected Akaike information criterion (AICc, Hurvich and Tsai 1989) for AR models using 0 to 15 AR terms, selecting the model with the lowest score. This procedure optimizes the tradeoff between a model’s goodness-of-fit and the number of parameters estimated (Akaike, 1974). Parameters were fit by maximizing the likelihood, using a state-space representation of the AR process to handle the missing values (Jones, 1980). Residuals were tested for difference from white noise at the 0.05 level using the Ljung-Box test (Ljung and Box, 1978). All analyses were run in R (R Development Core Team, 2014).

## 3. Results

### 3.1. Seasonal cycles

Weather and oceanography followed a clear annual cycle (Figure 2, Table 1). Northwesterly winds and offshore Ekman transport were strongest in early spring, co-occurring with low SSH and high fluorescence. The SSH anomaly was lowest in April and May. SST and SSH had the strongest seasonal patterns, with the seasonal models accounting for 39% and 50% of their overall variabilities (Table 1). Upwelling and fluorescence were more episodic, with only 13% and 14% of their variabilities explained by the seasonal models (Table 1).

**Figure 2:**
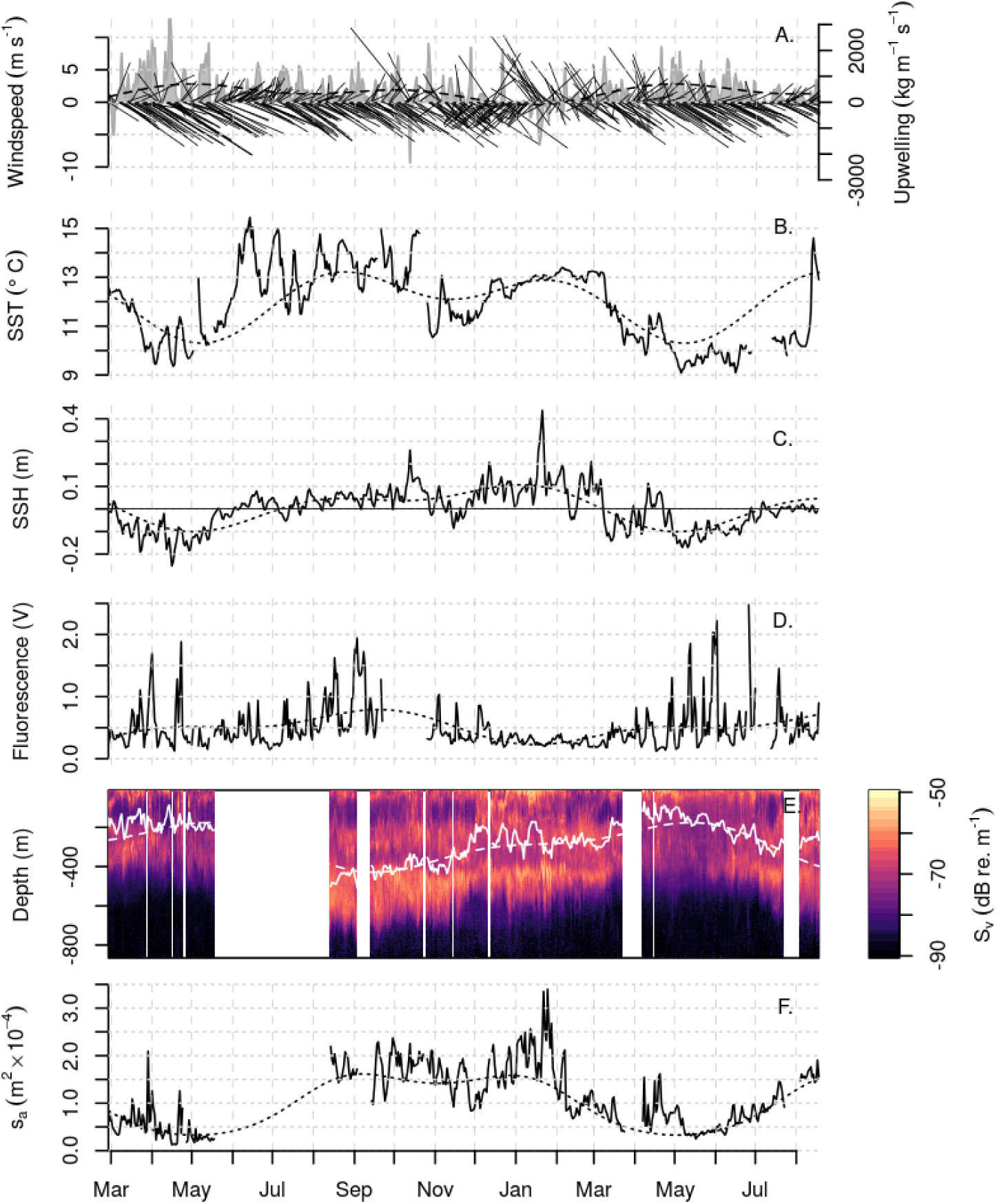
Oceanographic and acoustic time series from 27 February, 2009, to 18 August, 2010, with seasonal sinusoidal models (dotted lines). A) Wind vectors (magnitudes on left axis) and calculated upwelling (gray area, right axis). B) Sea-surface temperatures. C) Fluorescence. D) Sea-surface height. E) Daytime acoustic backscatter. White line shows vertical center of mass (CM). F) Depth-integrated area backscattering coefficient (*s*_*a*_).

**Table 1:** Summary of sinusoidal models fit to environmental and acoustic time series.

| Variable | Units | Mean | High | (date) | Low | (date) | $R^2$ |
| --- | --- | --- | --- | --- | --- | --- | --- |
| Upwelling | $\text{kg (m s)}^{-1}$ | 341.27 | 703.3 | Apr 30 | -97.5 | Jan 11 | 0.13 |
| SST | $^{\circ}\text{C}$ | 12.07 | 13.2 | Aug 24 | 10.3 | May 7 | 0.39 |
| Fluorescence | V | 0.51 | 0.8 | Sep 18 | 0.2 | Jan 16 | 0.14 |
| SSH | m | 0.02 | 0.1 | Jan 5 | -0.1 | May 2 | 0.50 |
| CM | m | -296.99 | -175.8 | May 12 | -428.7 | Sep 15 | 0.70 |
| Sa | dB | -40.26 | -37.9 | Sep 6 | -44.8 | May 3 | 0.72 |

The acoustic variables also had distinct seasonal cycles. Depth-integrated backscatter was lowest in May, coinciding with the coolest temperatures and highest fluorescence. It was highest in the fall and winter, with the seasonal model peaking in September at ‐37.5 dB, while the highest overall *S*_*a*_ values came during a short spike to ‐34.7 dB over several days in January 2010. During its spring minima, backscatter moved up in the water column, with its CM near 155 m depth. Backscatter was deepest in the water column in September, centered near 420 m. This change in the CM was due to the formation of a deep, thick, mostly non-migratory scattering layer between 400 and 700 m depth (Figure 2). The seasonal model for *S*_*a*_ accounted for 71% of its variability, and the seasonal model for the CM accounted for 74% (Table 1).

The seasonal cycle of backscatter was out of phase with that of upwelling and primary production (Table1). The correlation coefficient of *S*_*a*_ with SST was ‐0.64; with SSH it was 0.73 (Figure 3). The CM was negatively correlated with SST (*r* = –0.6) and SSH (*r* = –0.48). All other correlation coefficients were less than 0.3 (Figure 3).

**Figure 3:**
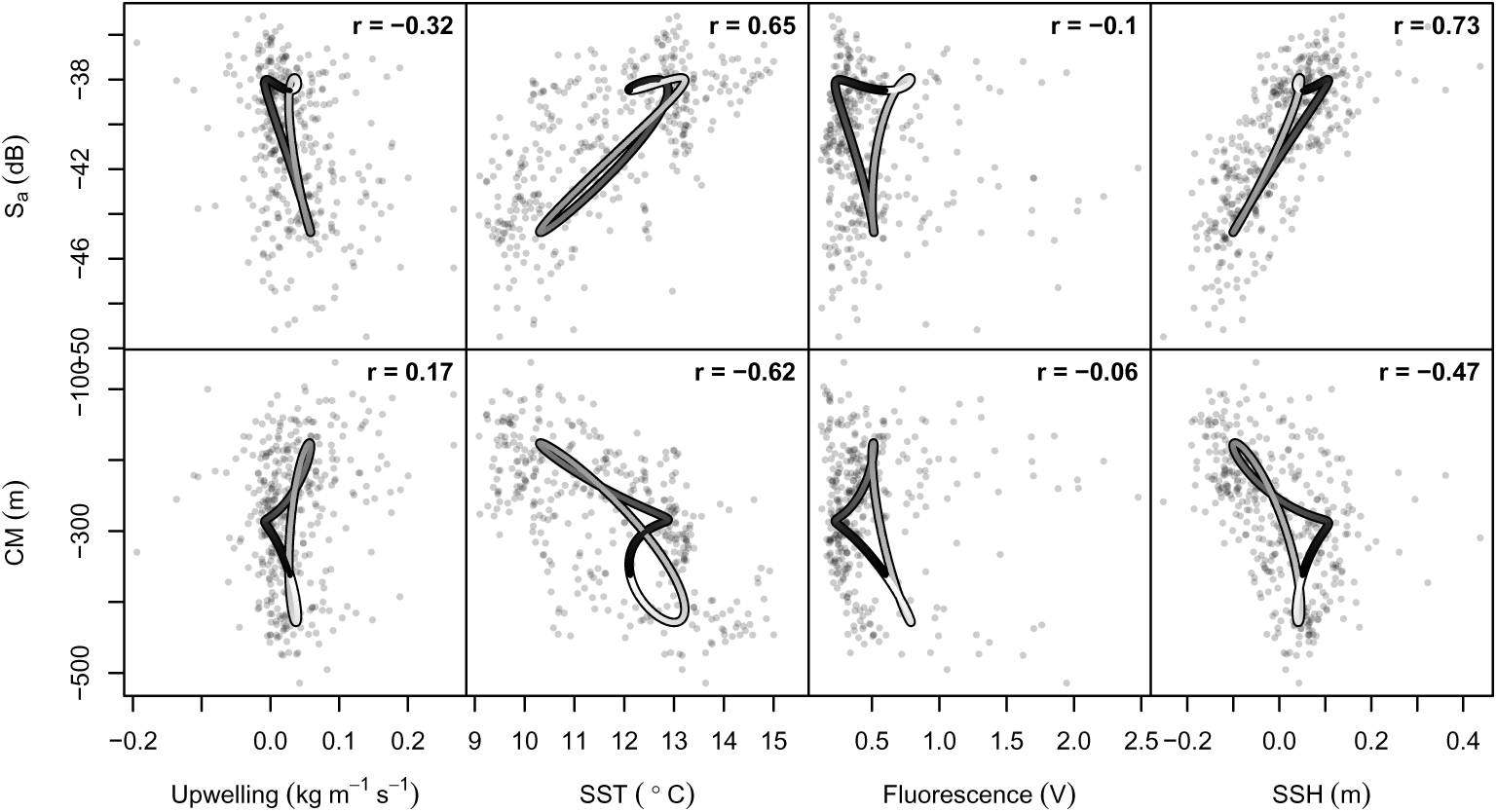
Pairwise relationships between oceanographic and acoustic time series at the seasonal time scale. Oceanographic variables, in columns from left to right, are upwelling Ekman transport, sea surface temperature (SST), chlorophyll fluorescence, and sea-surface height (SSH). Acoustic variables, in rows, are total depth-integrated backscatter (*s*_*a*_) and the acoustic center of mass (CM). Points show daily values of the raw time series, with Pearson’s product-moment correlation coefficient (*γ*) displayed in upper-right corner. Curves show the modeled seasonal cycles for each pair of variables. The curves’ color indicates day of the year, starting with black on 1 January and ending with white on 31 December. Because these models are sinusoidal, they must form loops, ending where they started. The closer a loop is to a straight line, the closer the cycles are to being perfectly in phase.

### 3.2. Sub-seasonal dynamics

At sub-seasonal time scales, fluorescence was negatively correlated with SST, with the highest correlation found at a lag of 3 days (Figure 4). Fluorescence displayed a characteristic scale of variability between 10 and 20 days, representing the average time between upwelling events and corresponding phytoplankton blooms. Sea level also showed semi-periodic fluctuations with a period of approximately 20 days. These fluctuations were correlated (*ρ* = –0.45) with alongshore wind stress at lags of 1 day (Figure 4).

**Figure 4:**
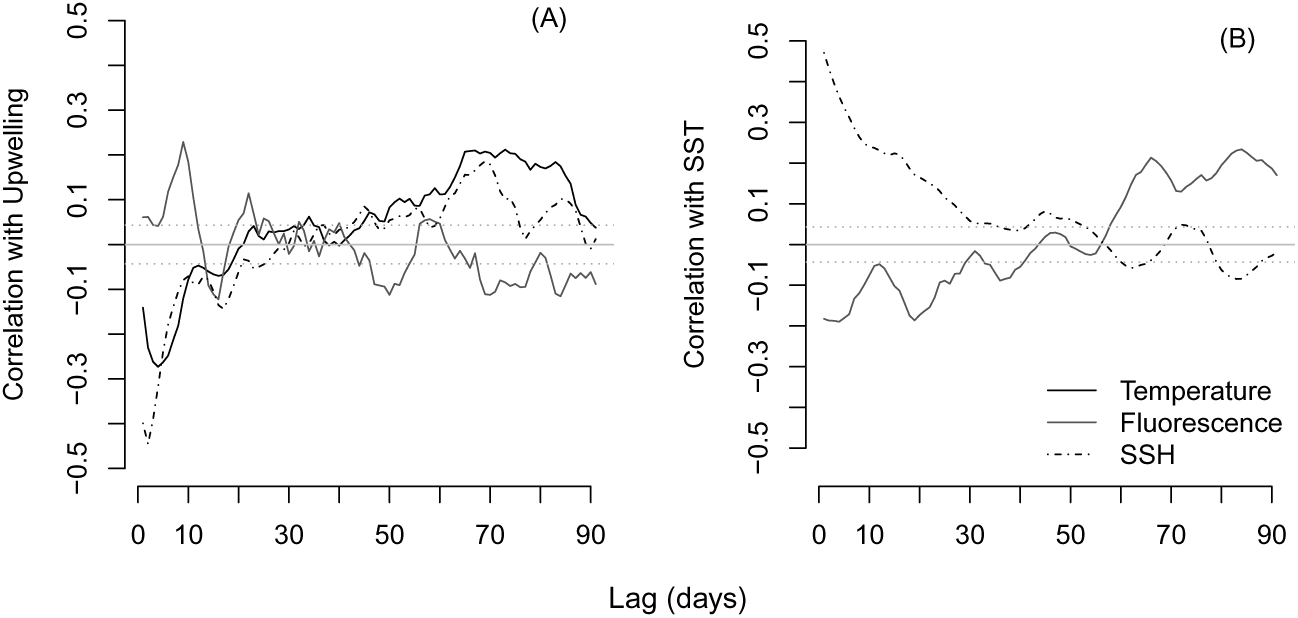
Cross-correlations between environmental series. A) Lagged correlations of sea-surface temperature, sea level, and fluorescence with alongshore wind stress. B) Lagged correlations of sea level and fluorescence with se-surface temperature. Dotted grey lines show significance at the 0.05 level for a white-noise null hypothesis.

The density of pelagic animals was also related to oceanographic variability at sub-seasonal time scales. Total backscatter had weak but significant negative correlations with indicators of upwelling (alongshore wind stress, below-average SST and above-average fluorescence) at lags less than 20 days (Figure 5). The strongest relationship between upwelling variables and animal distribution was found in the CM, which was negatively correlated with SST at all lags, with its minimum (*ρ* = ‐0.23) at 14-15-day lags. The CM was also negatively correlated with fluorescence at lags between 0 and 18 days, though not as strongly as with temperature. Taken together, these correlations indicate that the total animal abundance decreased slightly over one to three weeks following upwelling events, while moving up in the water column.

**Figure 5:**
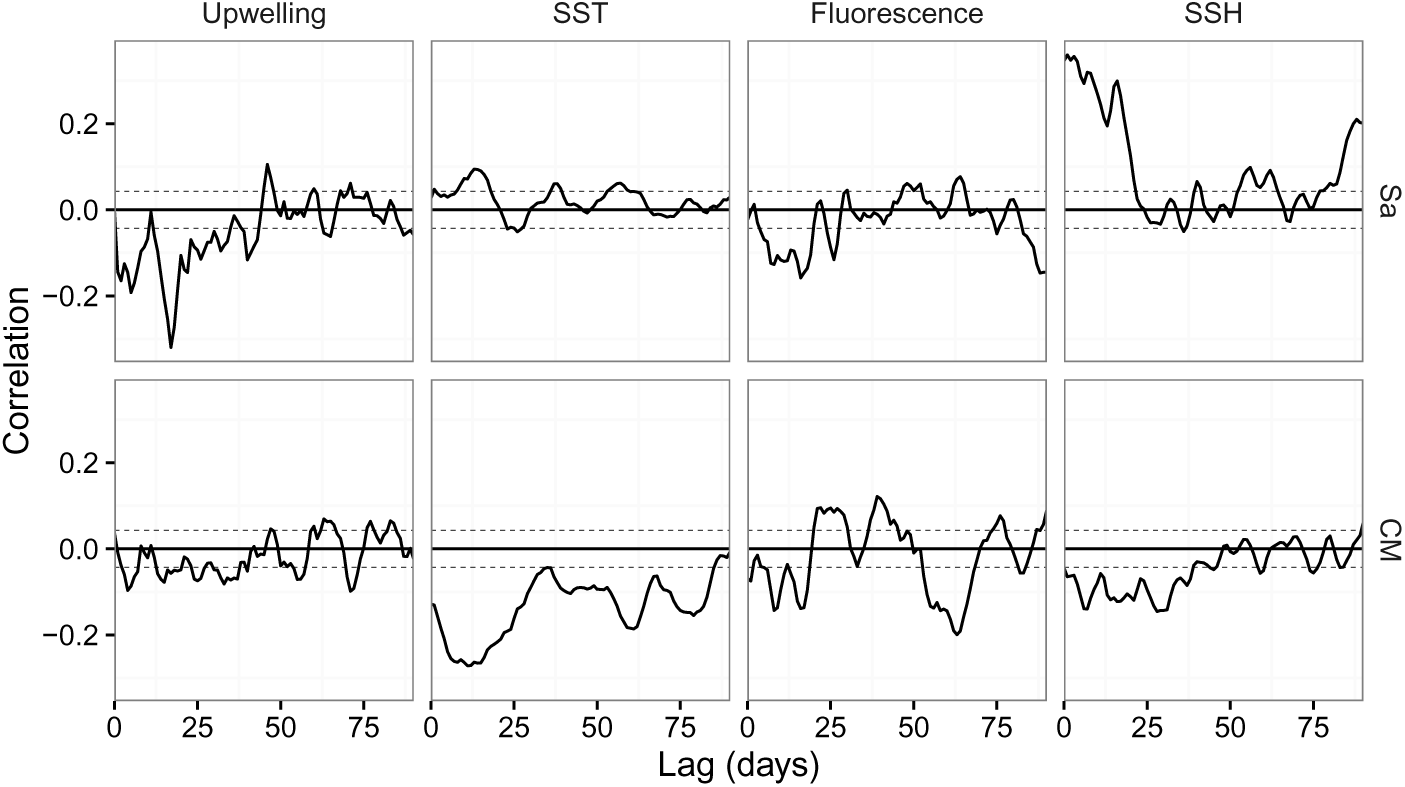
Cross-correlations between acoustic and environmental series. Plots show lagged correlation of total backscatter (*s*_*a*_, top row) and its mean location in the water column (the center of mass CM, bottom row) with alongshore wind stress, sea-surface temperature (SST), Fluorescence, and sea-surface height (SSH). Dotted lines show significance at the 0.05 level for a white-noise null hypothesis.

Of the environmental series examined, backscatter was best correlated with sea level. Cross-correlation of total backscatter with sea level was highest (*ρ* = 0.31) at a 0-day lag. The center of mass was significantly negatively correlated with sea level at lags from 0 to 38 days. Together, these correlations indicate that above-average sea levels were associated with increased backscatter deeper in the water column.

The correlation of backscatter to environmental variability also varied as a function of depth (Figure 6). Backscatter through the water column was negatively correlated with alongshore wind stress, indicating decreases in animal density immediately following upwelling winds (Figure 6A). At 40-75 day lags, the CCF became positive around 300 and 600 m, indicating that backscatter increased at these depths.

**Figure 6:**
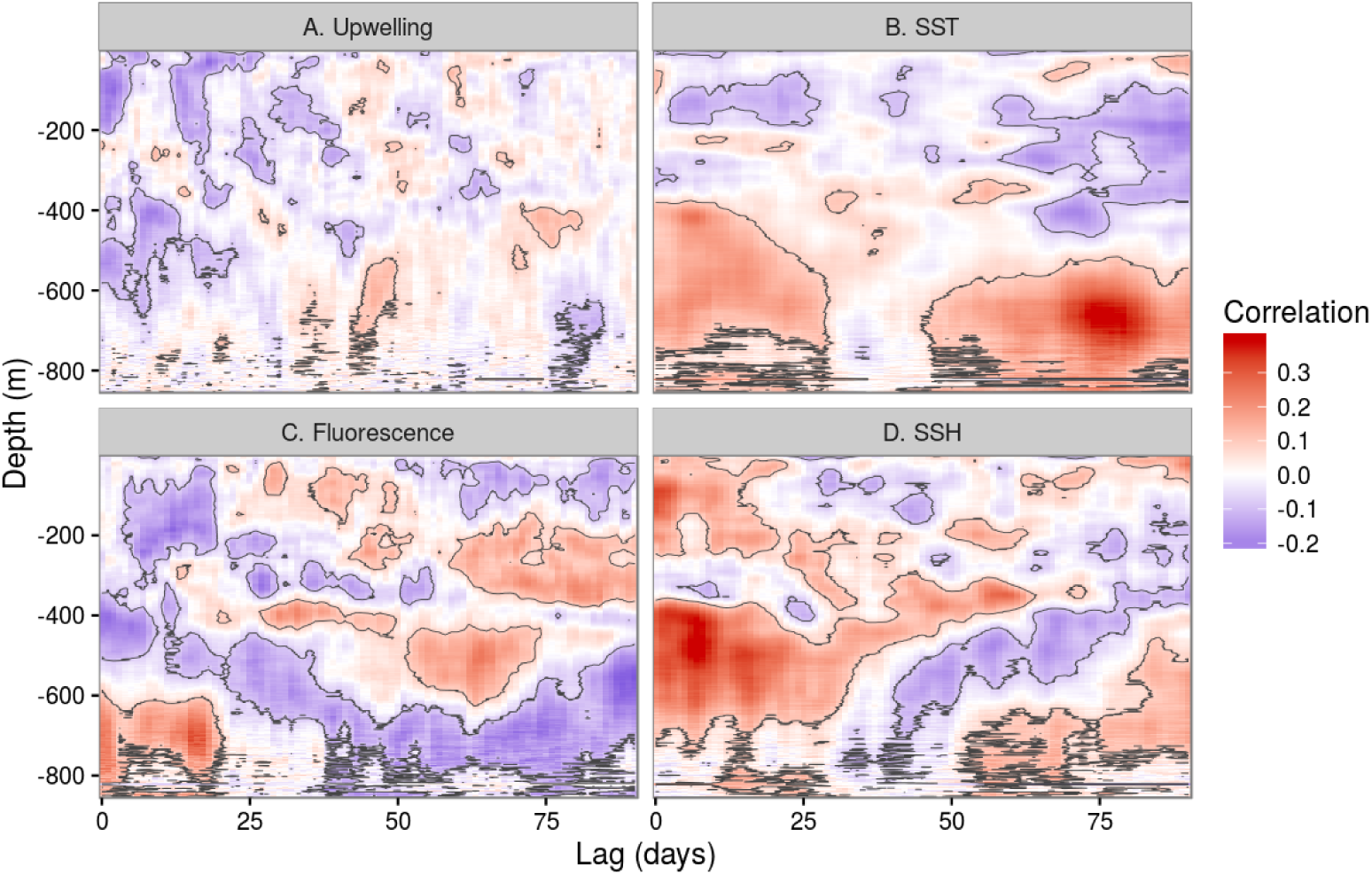
Correlations between environmental series and acoustic backscatter, as a function of lag (x-axis) and depth (y-axis). Red represents positive correlation, while blue represents negative correlation. Black contours enclose areas significantly different from zero at the 0.05 level, assuming a white-noise null hypothesis. Sub-figures show correlations between backscatter and A) alongshore (i.e., upwelling) wind stress, B) sea-surface temperature, C) surface fluorescence, and D) sea-surface height.

The effects of temperature on backscatter depended more on depth than on time lag. Backscatter above 200 m was negatively correlated with surface temperature at lags from two to almost 90 days. Conversely, backscatter at all depths below about 500 m was negatively correlated with temperature at all lags (Figure 6B). These results agreed with those for the CM, and indicate that decreases in surface temperature were associated with a long-lasting increase in animal density in the upper water column, and a corresponding decrease in density in the mesopelagic zone.

Backscatter was negatively correlated with fluorescence at lags from 0-20 days through most of the water column above 600 m, indicating a decrease in animal density following phytoplankton blooms (Figure 6C). The correlation became positive at 20-25 day lags near the surface. Below 600 m, this relationship appeared to be reversed. At increasing lags, positive correlations appeared at deeper depths, in an apparent propagation down the water column to about 300 m at a 60 day lag behind fluorescence. A similar downward movement of the fluorescence signal was apparent over approximately the same range of temporal lags, but deeper, beginning near 350 m and descending to about 550 m depth (Figure 6C).

For the regression model of micronekton in the upper 300 m of the water column, wind stress at a 5-day lag and sea level at a 0-day lag were significant (*p* < 0.05) predictors of backscatter above 300 m. The AICc procedure selected a model including these covariates and seven past values of backscatter (Table 2). The model’s one-step-ahead prediction errors were uncorrelated (all Ljung-Box p-values > 0.8), with an error variance of 1.57 (dB re. 1 m^2^ m^‒1^)^2^. In the simple linear regression with no autoregressive component, wind and sea level explained 8% of the variability. The addition of seven autoregressive terms improved the *R*^2^ value to 0.53.

**Table 2:** Parameter estimates, standard errors, and units for model of backscatter (dB re. 1 m^2^ m^‒2^) in the upper 300 m of the water column: 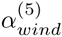 is the regression coefficient for wind stress at a 5-day lag, 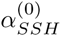 is the regression coefficient for sea-surface height at a 0-day lag, and *ϕ*^(*h*)^ is the autoregressive coefficient at lag *h* (in days).

| Parameter | $\alpha_{wind}^{(5)}$ | $\alpha_{SSH}^{(0)}$ | $\phi^{(1)}$ | $\phi^{(2)}$ | $\phi^{(3)}$ | $\phi^{(4)}$ | $\phi^{(5)}$ | $\phi^{(6)}$ | $\phi^{(7)}$ |
| --- | --- | --- | --- | --- | --- | --- | --- | --- | --- |
| Estimate | 7.336 | -8.170 | 0.657 | -0.055 | 0.120 | 0.025 | -0.018 | -0.200 | 0.252 |
| St. Error | 2.036 | 1.447 | 0.050 | 0.062 | 0.062 | 0.066 | 0.064 | 0.062 | 0.051 |
| Units | dB m<sup>-1</sup> | dB m<sup>-1</sup> | - | - | - | - | - | - | - |

### 4. Discussion

#### 4-1. Identity and biology of scattering species

Though it is not possible to determine species composition from single-frequency acoustic data without direct sampling, we can use knowledge of the backscattering properties of common zooplankton and fish (Stanton et al., 1996; Horne and Clay, 1998) and literature on the California Current (Barham, 1956; Kalish et al., 1986) to attribute most of the observed backscatter to mesopelagic micronekton.

Micronekton in the California Current are dominated by a relatively small number of species, including krill (*Euphausia pacifica* and *Thysanoessa spinifera*), *Sergestes similis*, a panaeid shrimp, and myctophid fishes (*Dia-phus theta, Stenobrachius leucopsarus*, and *Tarletonbeania crenularis*) (Phillips et al., 2009). All of these animals have been associated with sound-scattering layers (Barham, 1956; Kalish et al., 1986). Larger nekton are present as well, including macrourids (Yeh and Drazen, 2011), Pacific hake (*Merluc-cius productus*) and, in recent years, Humboldt squid (*Dosidicus gigas*, see Field et al. 2007), though the relative contribution of these species to overall biomass and backscatter is likely to be small compared to the smaller but much more abundant micronekton species. When the acoustic threshold on 30 randomly-selected echograms was raised from ‐90 dB (the level used in our analysis) to ‐58.5 dB (the level used for fisheries surveys of Pacific hake *Mallotus villosus*, Wilson and Guttormsen 1997), an average of 77% of the backscatter was eliminated. This difference suggests that the majority of backscatter is attributable to smaller animals.

Dense surface aggregations were sometimes present, especially from February through April, probably representing surface-schooling krill (*Euphausia pacifica* or *Thysanoessa spinifera*, Smith and Adams 1988), sardine (*Sardinops sagax*), or anchovy (*Engraulis mordax*, Cailliet et al. 1979). These aggregations make substantial contributions to water column biomass on the temporal scale of minutes as they pass through the acoustic beam, but do not affect trends at scales of days or longer.

Visual surveys from ROVs have shown that a substantial portion of Monterey Bay’s mesopelagic fauna is gelatinous (Robison, 2004; Robison et al., 2010). Though often considered weak acoustic targets relative to swimblad-dered fish, gelatinous animals may in fact make substantial contributions to backscatter (Colombo et al., 2003). The physonect siphonophore *Namomia bijuga* may be particularly important, since it is both abundant in Monterey Bay and posesses a gas-filled pneumatophore (Stanton et al., 1998; Warren, 2001). *Nanomia’s* seasonal cycle matches that of backscatter quite well, peaking in summer between 200 and 600 m depth (Robison et al., 1998). Siphonophores have long been recognized as potential contributors to DSLs (Barham, 1963), but they are probably even worse-represented in net catches than mesopelagic fish (Hamner et al., 1975; Kaartvedt et al., 2012).

#### 4.2. Seasonal cycles

The environmental time series showed typical seasonal cycles for Monterey Bay and the California Current. Northwesterly winds were highest in spring, along with the surface expression of cooler water and spikes in fluorescence, indicating upwelling and blooms of phytoplankton. Sea surface height at the coast was also lowest in spring, likely associated with upwelling-favorable winds pushing the surface layer offshore and lower the sea level near the coast.

At the same time, the acoustic center of mass moved up in the water column and overall backscatter decreased. This cycle agrees with previous measurements from an ADCP on the M1 mooring (Croll et al., 2005), and appears similar to recent measurments from the monsoon-driven upwelling system in the Arabian Sea (Wang et al., 2014). The upward movement of the CM may be explained by avoidance of a shoaling oxygen minimum zone (OMZ). An oxygen minimum zone is found between approximately 500 and 1000 m depth in Monterey Bay, and is associated with a decrease in animal density and changes in species assemblages (Lynn et al., 1982; Robison et al., 2010). Globally, DSLs are found higher in the water column where dissolved oxygen is lower (Klevjer et al., 2016). The OMZ in eastern boundary currents rises during upwelling, changing the distribution of micronekton and fish (Escribano et al., 2000; Chan et al., 2008; Bertrand et al., 2011). The decrease in overall backscatter may be attributed to offshore transport of animals in the upper water column.

Alternatively, these changes in the distribution of animals could be due to seasonal cycles of reproduction and population dynamics. The winter backscatter peak also appeared to agree with annual cycles of common micronekton in the California Current. *Euphausia pacifica*, the dominant krill species, spawn mostly in the spring, with adult abundance peaking in fall (Marinovic et al., 2002). Market squid (*Loligo opalescens*) are most abundant in Monterey Bay from April through July (Fields, 1965). *Sergestes similis* reproduce year-round with a springtime peak, and are most abundant over the continental slope during winter (Pearcy and Forss, 1969; Omori and Gluck, 1979). Myctophids in southern California were most abundant in winter (Paxton, 1967). With the exception of squid, these animals’ life-histories are consistent with the observed seasonal changes in backscatter, supporting our attribution of most of this acoustic energy to micronekton.

#### 4.3. Sub-seasonal dynamics

##### 4.3.1 Backscatter and sea level

At sub-seasonal time scales, sea surface height had the strongest and most immediate correlation with acoustic backscatter. This correlation was present through the water column, and suggests that advection is the most important physical processes affecting the density of micronekton on time scales of days to weeks.

Variation in coastal sea level is caused by several different processes, including atmospheric pressure, wind-driven upwelling, and coastally-trapped waves (Chelton and Enfield, 1986). Changes in atmospheric pressure force a static response in sea-surface height, which rises approximately 1 cm per 1 mbar drop in air pressure (the “inverted barometer” effect, Chelton and Enfield 1986). As described above, wind also affects sea level by pulling the surface layer away or pushing it towards the coast. Together, these two factors (measured at M1) explain 63% of the variability in our de-seasonalized sea level data.

Nearshore sea levels also vary with the passage of coastally trapped Kelvin waves, which propagate northward along the California coast (Enfield and Allen, 1980; Lyman and Johnson, 2008). Marinovic et al. (2002) found evidence that Kelvin waves moved southern zooplankton species into Monterey Bay during the 1997-1998 El Niño. Clarke and Dottori (2008) also found that aggregate zooplankton biomass in the southern California Current was correlated with sea level at San Diego with a two-month lag. They attributed this correlation to enhanced upwelling, primary production, and zooplankton population growth behind the waves, where sea level is lowest and the thermocline is shallowest.

In contrast to Clarke and Dottori’s (2008) results, the correlations observed in this study occurred at a scale of days to weeks, with no lag. The near-immediate response of micronekton to sea level change, the consistency of this response through the water column, and the days-to-weeks-scale correlation all suggest passive aggregation by fluid motion. This result aligns with the analysis in Urmy et al. (2012), which showed that the power spectrum of the backscatter time series from DEIMOS was similar to that expected for the velocity of a turbulent fluid.

A dramatic example of an immediate response occurred from 16-31 January 2010, when SSH and backscatter both spiked to their 18-month highs within the same 3-day period (Figure 2, Figure 7). On January 18, 19, and 20, strong southeasterly winds caused onshore Ekman transport (Figure 7B). Combined with low low atmospheric pressure (986.6 mbar, measured at M1) this transport led to the highest sea level of the 18-month series, increasing from 0.1 to 0.5 m above normal (Figure 7C). Shortly before the peak SSH on 21 January, depth-integrated backscatter began to rise, increasing four-fold from its minimum on 20 January (Figure 7D). This increase was driven by an abrupt thickening of the deep scattering layer centered around 400 m depth (Figure 7A). SSH declined over the next two days, rising slightly again on 25-26 January with another, weaker, southeasterly wind event. Backscatter fell on 24 Janurary, then spiked again to its 18-month high (0.46 × 10^‒^4 m^2^) on 25 January before declining back to its previous level (1.2 × 10^‒^4 m^2^) over the next five days.

**Figure 7:**
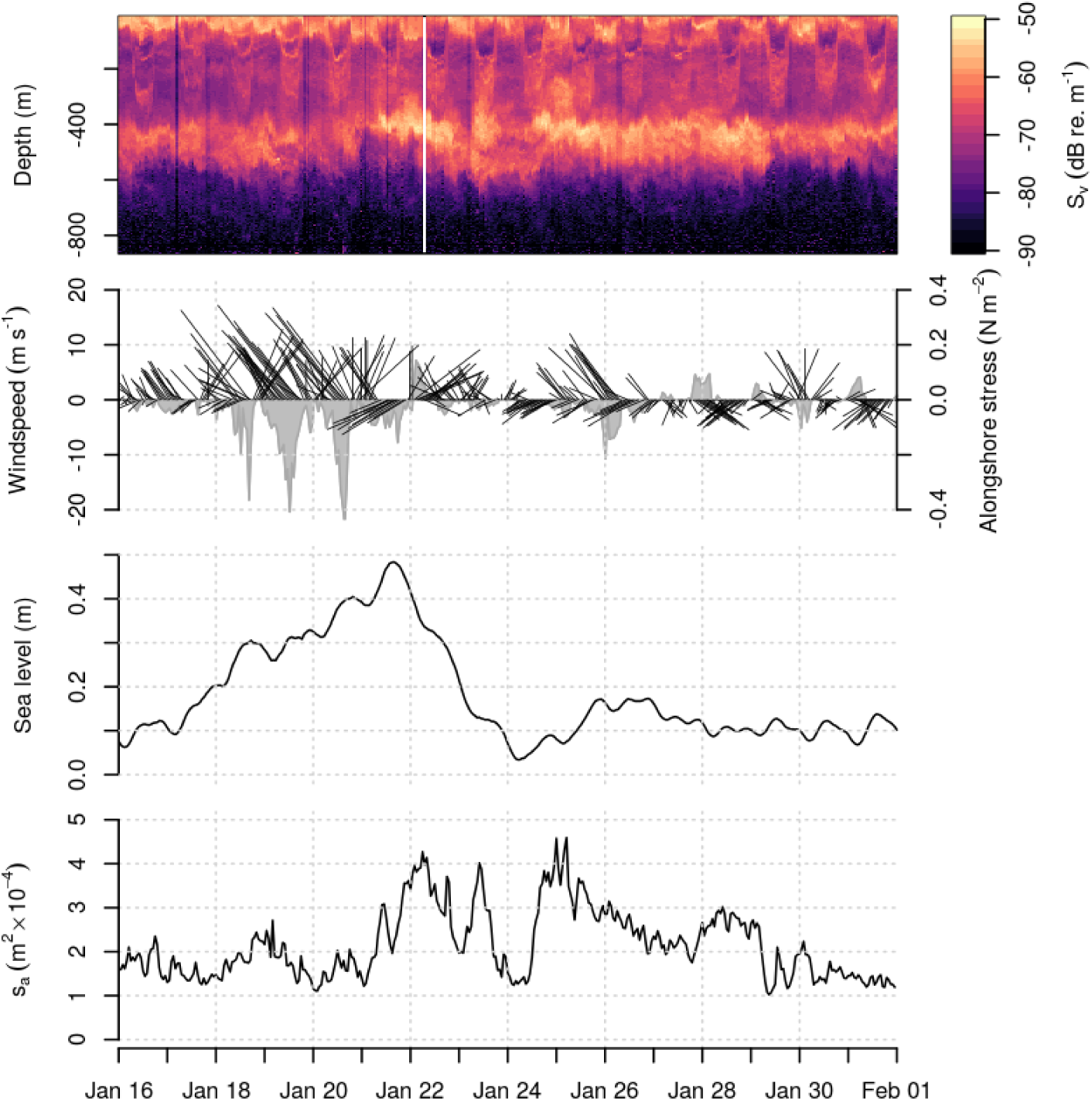
Hourly wind, sea level, and backscatter from 16-30 January, 2010. A) Echogram, showing volume backscattering strength (color) as a function of depth (y-axis) and time (x-axis). B) Wind direction and alongshore wind stress, as in Figure 2. C) Low-pass filtered Monterey sea level. D) Depth-integrated backscatter, approximately proportional to water column biomass. Strong southeasterly winds from 18-20 January precede a rise in sea level and total backscatter the next day.

Several mechanisms could be proposed to explain the correlation between SSH and backscatter. Onshore Ekman transport could collect zooplankton and micronekton in the surface layers against the coast, but would not extend to the mesopelagic zone, where sea level and micronekton were also correlated. Downwelling resulting from onshore Ekman transport could perhaps carry animals from the surface to deeper water, but Ekman upwelling/downwelling velocities in the California Current are on the order of 10 m d^‒1^ (Huyer, 1983; Miinchow, 2000), and do not explain the sudden thickening of a biological layer centered at 400 m depth. Alongshore advection of a pre-existing aggregation is another possibility. Flagg et al. (1994), in a long-term acoustic Doppler current profiler deployment in the Mid-Atlantic Bight, observed similar abrupt (i.e. day-scale) increases in backscatter associated with reversals in alongshore currents.

Offshore, mesoscale eddies and jets can aggregate zooplankton (Huntley et al., 2000) and alter deep scattering layers (Kloser et al., 2009; Godø et al., 2012; Fennell and Rose, 2015). Both Godø et al. (2012) and Fennell and Rose (2015) found the DSL thickened under anticyclonic eddies, where the SSH is anomalously high and the isopycnals are deflected downwards. Anticyclonic eddies are associated with downwelling and low primary production, suggesting the thickened DSL is due to physical aggregation. Indeed, boundaries of the DSL appear to closely track the isopycnals, further suggesting passive transport (Godø et al., 2012).

Echograms of thickened DSLs under anticyclonic eddies (Godø et al., 2012; Fennell and Rose, 2015) appear qualitatively similar to Figure 7. While we did not measure hydrographic profiles to accompany our SSH data, isostatic balance requires a downward deflection in the isopycnals accompanying an upward deflection of the sea surface. If an anticyclonic eddy were to impinge on the shelf, it could potentially bring zooplankton and micronekton with it. A coastally-trapped Kelvin wave would have similar effects, since it it is, in a sense, a geostrophic eddy with a coastline running through its center (Gill and Clarke, 1974; Wang and Mooers, 1976).

##### 4.3.2. Backscatter and upwelling

The response of micronekton to upwelling events was less pronounced than the response to sea level, but still measurable. At time lags less than one month following low SST, backscatter decreased in the mesopelagic and increased in the epipelagic, while decreasing slightly overall. This is consistent with upward animal movement, to avoid the shoaling of the OMZ, followed by transport offshore with the surface layer. An alternate possibility is that increased primary production near the surface shaded the water column below. Similar changes in downwelling irradiance can induce dramatic changes in the distribution of zooplankton (Frank and Widder, 2002).

The negative correlation between backscatter and fluorescence at lags less than 20 days is related to correlations of backscatter with sea level and temperature, since upwelling events that precede phytoplankton blooms are also associated with decreases in sea level and temperature. Decreased micronekton abundances following phytoplankton blooms are interpreted as a product of these physical processes, rather than a biological response to phytoplankton production. We cannot explain the positive correlation between backscatter and fluorescence below 600 m (Figure 6C) from 0-20 day lags. However, *s*_*v*_ at these depths was 20-40 dB (i.e., 2-4 orders of magnitude) lower than in the main scattering layers, so the biological significance of these correlations is expected to be minor.

At time lags greater than 20 days, there appeared to be a depth-dependent response of micronekton to increased productivity at the surface. Positive correlations of backscatter with fluorescence peaked at the surface at approximately 20-day lags, and propagated downward to 300 m within another 20-30 days, translating to a speed of 10-15 m d^‒1^. This rate is faster than sinking phytoplankton (up to 1.69 m d^‒1^, Bienfang 1980), and slower than the fecal pellets of krill (126-862 m d^‒1^, Fowler and Small 1972), midwater fish (1028 m d^‒1^, Robison and Bailey 1981), or larvacean houses (800 m d^‒1^, Robison et al. 2005). It is slower than sinking aggregates at the end of the North Atlantic spring bloom (75 m d^‒1^, Briggs et al. 2011), but agrees fairly well with sinking rates of marine snow measured in Monterey Bay (16.29 to 25.46 m d^‒1^ depending on particle size, Pilskaln et al. 1998).

The timing of these depth-dependent cross-correlations is suggestive of a near-surface pulse of secondary production in zooplankton following up-welling events, which then propagates down the water column as marine aggregates. Detritus and marine snow are weak scatterers, but not acoustically invisible. Sinking pulses of krill fecal pellets can be resolved at 200 kHz (Røstad and Kaartvedt, 2013), and it is possible that some aggregates contain gas bubbles, which would make them “visible” at 38 kHz (Opdal et al., 2008). Sinking aggregates can also support dense communities of zooplankton with benthic-like morphology and feeding behavior (Steinberg et al., 1994), which would also increase their acoustic cross-section. Alternatively, free-swimming animals could track pulses of sinking detritus down the water column, since these particles represent a valuable food resource in the deep ocean (McClain, 2010).

##### 4.3.3. Strength of sub-seasonal coupling

While statistically significant relationships were detected between environmental and acoustic time series at sub-seasonal time scales, none of these relationships were particularly strong. The maximum correlation coefficient (between sea level and backscatter) was only 0.31, and in the lagged-regression model, far more variance was explained by autocorrelation than by any of the explanatory variables. Several aspects of the study’s design may have masked physical-biological relationships. DEIMOS and Ml were separated by approximately 15 km, and even though physical measurements were correlated across this distance, the separation between the two sensors culd have obscured relationships. Similarly, DEIMOS’s acoustic beam spanned only 115 m horizontally at its widest point, which might not be an appropriate scale to observe the biological-physical coupling (cf. Schneider and Piatt 1986).

The response of micronekton to upwelling and other environmental variability could simply be weak at sub-seasonal time scales. Population dynamics, somatic growth, and kinematics (i.e., passive or active aggregation of animals in productive areas) could all link environmental variability to changes in biomass and backscatter. While there is evidence that some micronekton cue reproduction on upwelling events (Dorman et al., 2005), the productivity-population relationship is not expected to be strong. Mid-trophic-level micronekton must wait for smaller zooplankton to increase their populations first. In effect, each successive trophic level acts as a low-pass filter, smoothing out variability at scales shorter than its generation time.

Somatic growth could also account for increases in backscatter. As an example, Moku et al. (2001) estimated growth rates for juvenile *Diaphus theta*, the most common myctophid in DSLs in the California Current (Barham, 1956; Kalish et al., 1986). Combined with a target-strength/length relationship from Yasuma et al. (2003), we can predict target strength (dB re. 1 m^2^) as a function of age in days (t) as *TS* = 11.83 log_10_(0.354 ‫ 0.0129t) – 63.53. This equation implies that cohort of juveniles could increase its scattering cross-section 36% in one month. On the other hand, some mesopelagic fishes have swim bladders that deflate or fill with wax esters as they reach adulthood, leading to *decreases* in target strength (Yasuma et al., 2010). In any case, somatic growth is not expected to respond quickly to individual pulses of food.

Aggregation through active swimming also seems unlikely. Though myctophids can swim up to 8 cm s^‒1^, translating to 6.9 km d^‒1^ (Kaartvedt et al., 2009), they have little reason to travel horizontally. Consider an average *Diaphus theta* weighing 0.4 g (Sassa et al., 2002), which must eat 1-6% of its weight in food each day (Kosenok et al., 2006; Moku et al., 2000). If, during several hours of nighttime foraging at the surface, it capures a single medium-sized krill (10-40 mg for *Euphausia pacifica*, Wilson et al. 2009), its energy needs will be met. Directed horizontal swimming is therefore probably not energetically favorable for vertically migrating micronekton Indeed, most myctophids tracked in a deep fjord drifted passively at depth (Kaartvedt et al., 2009).

### 5. Conclusion

Our results illustrate a complex relationship between the variability of animal biomass density and variability in their environment. Biomass density at any location varies as animals swim, drift with currents, grow, interact with other species, and increase or decline in population. Quantifying the relative importance of these processes depends on the spatial and temporal scales over which they occur, as well as measurement resolution (Horne and Schneider, 1994).

The seasonal cycle of backscatter was negatively correlated and almost exactly out of phase with that of upwelling. These changes were mostly due to a deep, non-migrating scattering layer centered near 500 m depth, which disappeared during the spring upwelling season and thickened to its maximum during the fall and winter oceanic period.

Correlations between oceanography and the distribution of animals were present at sub-seasonal time scales, though they were weaker than those observed at seasonal scales. During upwelling events, the overall abundance of animals decreased and moved upwards, consistent with vertical swimming to avoid the shoaling OMZ and movement offshore with the Ekman layer. Approximately three weeks after upwelling events, backscatter increased in the surface layer, suggesting reproduction of small or medium-sized zooplankton. The upwelling signal then appeared to propagate down the water column at rates similar to those measured for sinking marine aggregates. The strongest physical-biological correlations at short time scales appeared to be driven by passive aggregation of micronekton by fluid motion.

Variability in animal density influenced by physical processes is distributed across a wide range of temporal scales. High-resolution, temporally-indexed observations of animal density allow variability in animal densities to be measured and compared to other biological and physical processes at temporal scales not possible when sampled using mobile platforms. Ultimately, stationary acoustic instruments could be used to augment fisheries and ecosystem assessments by adding independent, temporal indices of population abundance, flux, and variance.

Analyses used here were correlative and linear simplifications of a complex, dynamic pelagic ecosystem. More realistic dynamic models for the changing distribution of animals, explicitly based on physics and biology, would ultimately be more appropriate and informative. While we currently lack the detailed knowledge necessary to build such models, acoustic observing systems like DEIMOS can identify patterns in the underwater “landscape” and highlight complementary measurements necessary to interpret the acoustic records. In the present work, we have attempted to start this process.

## 6. Acknowledgements

We thank Kongsberg Maritime for the loan of the echosounder used in this study. David Barbee and Dick Kreisberg were instrumental in the design and assembly of DEIMOS. We also thank MBARI for hosting DEIMOS at the MARS observatory, particularly the crew and pilots of the RV *Point Lobos* and ROV *Ventana*. This paper owes much improvement to the helpful comments of two anonymous reviewers. Funding was provided by the University of Washington School of Aquatic and Fishery Sciences.

## References

Akaike, H., 1974. A new look at the statistical model identification. IEEE Transactions on Automatic Control 19 (6), 716–723.

Bakun, A., 1973. Coastal upwelling indices: West Coast of North America 1946–95. Tech. Rep. NMFS SSRF-693, U.S. Department of Commerce.

Barham, E. G., 1956. The ecology of sound scattering layers in the Monterey Bay area, California. Ph.D. thesis, Stanford University.

Barham, E. G., 1963. Siphonophores and the deep scattering layer. Science 140 (3568), 826–828.

Benoit-Bird, K. J.., Au, W. W. L., 2002. Energy: Converting from acoustic to biological resource units. The Journal of the Acoustical Society of America 111, 2070–2075.

Bertrand, A., Chaigneau, A., Peraltilla, S., Ledesma, J., Graco, M., Monetti, F., Chavez, F. P., 2011. Oxygen: a fundamental property regulating pelagic ecosystem structure in the coastal southeastern tropical Pacific. PloS one 6 (12), e29558.

Bienfang, P. K., 1980. Phytoplankton sinking rates in oligotrophic waters off Hawaii, USA. Marine Biology 61 (1), 69–77.

Block, B. A., Jonsen, I. D., Jorgensen, S. J., Winship, A. J., Shaffer, S. A., Bograd, S. J., Hazen, E. L., Foley, D. G., Breed, G. A., Harrison, A. L., Ganong, J. E., Swithenbank, A., Castleton, M., Dewar, H., Mate, B. R., Shillinger, G. L., Schaefer, K. M., Benson, S. R., Weise, M. J., Henry, R. W., Costa, D. P., 2011. Tracking apex marine predator movements in a dynamic ocean. Nature 475, 86–90.

Boersch-Supan, P. H.., Rogers, A. D., Brierley, A. S., 2015. The distribution of pelagic sound scattering layers across the southwest Indian Ocean. Deep Sea Research Part II: Topical Studies in Oceanography.

Bolin, R. L., Abbot, D. P., 1963. Studies on the marine climate and phytoplankton of the central coastal area of California, 1954-1960. CalCOFI Reports 9, 23–45.

Briggs, N., Perry, M. J., Cetinic, I., Lee, C., D’Asaro, E., Gray, A. M., Rehm, E., 2011. High-resolution observations of aggregate flux during a sub-polar North Atlantic spring bloom. Deep-Sea Research Part I: Oceanographic Research Papers 58 (10), 1031–1039.

Brinton, E., Townsend, A., 2003. Decadal variability in abundances of the dominant euphausiid species in southern sectors of the California Current. Deep-Sea Research II 50 (14-16), 2449–2472.

Brockwell, P. J., Davis, R. A., 2002. Introduction to Time Series and Forecasting, 2nd Edition. Springer-Verlag.

Brodeur, R., Yamamura, O., 2005. Micronekton of the North Pacific. PICES Scientific Report (30).

Cailliet, G. M., Karpov, K. A., Ambrose, D. A., 1979. Pelagic assemblages as determined from purse seine and large midwater trawl catches in Monterey Bay and their affinities with the market squid, Loligo opalescens. CalCOFI Reports 20, 21–30.

Chan, F., Barth, J. A., Lubchenco, J., Kirincich, A., Weeks, H., Peterson, W. T., Menge, B. A., 2008. Emergence of anoxia in the California Current Large Marine Ecosystem. Science 319, 920.

Chavez, F. P., Pennington, J. T., Herlien, R., Jannasch, H., Thurmond, G., Friederich, G. E., 1997. Moorings and drifters for real-time interdisciplinary oceanography. Journal of Atmospheric and Oceanic Technology 14 (1991), 1199–1211.

Chelton, D. B., Enfield, D. B., 1986. Ocean signals in tide guage records. Journal of Geophysical Research 91 (B9), 9081–9098.

Clarke, A. J., Dottori, M., 2008. Planetary wave propagation off California and its effect on zooplankton. Journal of Physical Oceanography 38 (3), 702–714.

Colombo, G. A., Mianzan, H., Madirolas, A., 2003. Acoustic characterization of gelatinous-plankton aggregations: four case studies from the Argentine continental shelf. ICES Journal of Marine Science 3139 (3), 1352–1360.

Croll, D. A., Marinovic, B. B., Benson, S. R., Chavez, F. P., 2005. From wind to whales: trophic links in a coastal upwelling system. Marine Ecology Progress Series 289, 117–130.

De Robertis, A., Higginbottom, I., 2007. A post-processing technique to estimate the signal-to-noise ratio and remove echosounder background noise. ICES Journal of Marine Science 64 (6), 1282–1291.

Demer, D. A., Conti, S. G., 2003. Validation of the stochastic distorted-wave Born approximation model with broad bandwidth total target strength measurements of Antarctic krill. ICES Journal of Marine Science 60 (3), 625–635.

Dietz, R. S., 1948. Deep scattering layer in the Pacific and Antarctic Oceans. Journal of Marine Research 7 (3), 430–442.

Dorman, J. G., Bollens, S. M., Slaughter, A. M., 2005. Population biology of euphausiids off northern California and effects of short time-scale wind events on Euphausia pacifica. Marine Ecology Progress Series 288, 183–198.

Enfield, D. B., Allen, J. S., 1980. On the structure and dynamics of monthly mean sea level anomalies along the Pacific coast of North and South America. Journal of Physical Oceanography 10, 557–578.

Escribano, R., Marin, V. H., Irribarren, C., 2000. Distribution of Euphausia mucronata at the upwelling area of Peninsula Mejillones, northern Chile: the influence of the oxygen minimum layer. Scientia Marina 64 (1), 69–77.

Fennell, S., Rose, G., 2015. Oceanographic influences on Deep Scattering Layers across the North Atlantic. Deep Sea Research Part I: Oceanographic Research Papers 105, 132–141.

Field, J. C., Baltz, K., Phillips, A. J., Walker, W. A., 2007. Range expansion and trophic interactions of the jumbo squid, Dosidicus gigas, in the California Current. CalCOFI Reports 48, 131–146.

Fields, W. G., 1965. The Structure, Development, Food Relations, Reproduction, and Life History of the Squid Loligo opalescens Berry. Fishery Bulletin 131, 108 pp.

Flagg, C., Wirick, C., Smith, S., 1994. The interaction of phytoplankton, zooplankton and currents from 15 months of continuous data in the Mid-Atlantic Bight. Deep-Sea Research II 41 (2-3), 411–435.

Foote, K. G., 1983. Linearity of fisheries acoustics, with addition theorems. The Journal of the Acoustical Society of America 73 (6), 1932–1940.

Foote, K. G., Knudsen, H. P., Vestnes, G., 1987. Calibration of acoustic instruments for fish density estimation: a practical guide. Tech. Rep. 144, International Council for the Exploration of the Sea, Copenhagen.

Fowler, S. W., Small, L. F., 1972. Sinking rates of euphausiid fecal pellets. Limnology and Oceanography 17 (2), 293–296.

Frank, T., Widder, E., 2002. Effects of a decrease in downwelling irradiance on the daytime vertical distribution patterns of zooplankton and micronekton. Marine Biology 140 (6), 1181–1193.

Gill, A. E., Clarke, A. J., 1974. Wind-induced upwelling, coastal currents and sea-level changes. Deep-Sea Research and Oceanographic Abstracts 21 (5), 325–345.

Godø, O.-R., Samuelsen, A., Macaulay, G. J., Patel, R., Hjøllo, S. S., Horne, J. K., Kaartvedt, S., Johannessen, J. A., 2012. Mesoscale Eddies Are Oases for Higher Trophic Marine Life. PLoS ONE 7 (1), e30161.

Graham, W. M., Field, J., Potts, D., 1992. Persistent “upwelling shadows” and their influence on zooplankton distributions. Marine Biology 114, 561–570.

Haeckel, E. H. P. A., 1890. Plankton-studien: Vergleichende untersuchungen uber die bedeutung und zusammensetzung der pelagischen fauna und flora. Verlag von Gustav Fischer, Jena.

Hamner, W. M., Madin, L. P., Alldredge, a. L., Gilmer, R. W., Hamner, P. P., 1975. Underwater observations of gelatinous zooplankton: Sampling problems, feeding biology, and behavior. Limnology and Oceanography 20 (6), 907–917.

Hays, G. C., 2003. A review of the adaptive significance and ecosystem consequences of zooplankton diel vertical migrations. Hydrobiologia 503, 163–170.

Horne, J. K., Clay, C. S., 1998. Sonar systems and aquatic organisms: matching equipment and model parameters. Canadian Journal of Fisheries and Aquatic Sciences 55 (5), 1296–1306.

Horne, J. K., Schneider, D. C., 1994. Analysis of scale-dependent processes with dimensionless ratios. Oikos 70 (2), 201–211.

Horne, J. K., Urmy, S. S., Barbee, D. H., 2010. Using sonar to describe temporal patterns of oceanic organisms from the MARS Observatory. Oceans 2010 MTS/IEEE Seattle, 1–7.

Huntley, M. E., Gonzalez, A., Zhu, Y., Zhou, M., Irigoien, X., 2000. Zooplankton dynamics in a mesoscale eddy-jet system off California. Marine Ecology Progress Series 201, 165–178.

Hurvich, C. M., Tsai, C.-L., 1989. Regression and time series model selection in small samples. Biometrika 76 (2), 297–307.

Huyer, A., 1983. Coastal upwelling in the California Current System. Progress in Oceanography 12 (3), 259–284.

Irigoien, X., Klevjer, T. A., Røstad, A., Martinez, U., Boyra, G., Acuna, J. L., Bode, A., Echevarria, F., Gonzalez-Gordillo, J. I.., Hernandez-Leon, S.., Agusti, S., Aksnes, D. L., Duarte, C. M., Kaartvedt, S., 2014. Large mesopelagic fishes biomass and trophic efficiency in the open ocean. Nature Communications 5 (May 2013), 3271.

Jones, R. H., 1980. Maximum likelihood fitting of ARMA models to time series with missing observations. Technometrics 22 (3), 389–395.

Kaartvedt, S., Røstad, A., Klevjer, T. A., Staby, A., 2009. Use of bottom-mounted echo sounders in exploring behavior of mesopelagic fishes. Marine Ecology Progress Series 395, 109–118.

Kaartvedt, S., Staby, A., Aksnes, D. L., 2012. Efficient trawl avoidance by mesopelagic fishes causes large underestimation of their biomass. Marine Ecology Progress Series 456, 1–6.

Kalish, J. M., Greenlaw, C. F., Pearcy, W. G., Holliday, D. V., 1986. The biological and acoustical structure of sound scattering layers off Oregon. Deep-Sea Research I 33 (5), 631–653.

Keister, J. E., Strub, P. T., 2008. Spatial and interannual variability in mesoscale circulation in the northern California Current System. Journal of Geophysical Research 113 (C4), C04015.

Kirk, J. T. O., 1994. Light and Photosynthesis in Aquatic Ecosystems. Cambridge University Press.

Klevjer, T. A., Irigoien, X., Røstad, A., Fraile-Nuez, E.., Benitez-Barrios, V. M.., Kaartvedt, S., 2016. Large scale patterns in vertical distribution and behaviour of mesopelagic scattering layers. Scientific Reports 6 (19873), 111.

Kloser, R., Ryan, T., Young, J., Lewis, M., 2009. Acoustic observations of micronekton fish on the scale of an ocean basin: potential and challenges. ICES Journal of Marine Science 66 (6), 998–1006.

Kosenok, N. S., Chuchukalo, V. I., Savinykh, V. F., 2006. The characteristics of feeding of Diaphus theta (Myctophidae) in the northwestern part of the Pacific Ocean in the Summer-Autumn period. J. Ichthyol. 46 (8), 606–612.

Legaard, K. R., Thomas, A. C., 2007. Spatial patterns of intraseasonal variability of chlorophyll and sea surface temperature in the California Current. Journal of Geophysical Research 112 (C9), C09006.

Ljung, G. M., Box, G. E. P., 1978. On a measure of lack of fit in time series models. Biometrika 65 (2), 297–303.

Lyman, J. M., Johnson, G. C., 2008. Equatorial Kelvin wave influences may reach the Bering Sea during 2002 to 2005. Geophysical Research Letters 35 (14), L14607.

Lynn, R. J., Bliss, K. A., Eber, L. E., 1982. Distributions of seasonal mean temperature, salinity, sigma-t, stability, dynamic height, oxygen, and oxygen saturation in the California Current, 1950-1978. CalCOFI Atlas 30, 432.

MacLennan, D. N., Fernandes, P. G., Dalen, J., 2002. A consistent approach to definitions and symbols in fisheries acoustics. ICES Journal of Marine Science 59 (2), 365–369.

Marinovic, B. B., Croll, D. A., Gong, N., Benson, S. R., Chavez, F. P., 2002. Effects of the 1997-1999 El Nino and La Nina events on zooplankton abundance and euphausiid community composition within the Monterey Bay coastal upwelling system. Progress in Oceanography 54, 265–277.

McClain, C., 2010. An empire lacking food. American Scientist 98 (6), 470.

McGowan, J. A., Bograd, S. J., Lynn, R. J., Miller, A. J., 2003. The biological response to the 1977 regime shift in the California Current. Deep-Sea Research II 50 (14-16), 2567–2582.

McGowan, J. A., Chelton, D. B., Conversi, A., 1996. Plankton patterns, climate, and change in the California Current. CalCOFI Reports 37, 4568.

Micheli, F., Cottingham, K. L., Bascompte, J., Eckert, G. L., Fischer, J. M., Keitt, T. H., Kendall, B. E., Klug, J. L., Rusak, J. A., 1999. The dual nature of community variability. Oikos 85 (1), 161–169.

Moku, M., Ishimaru, K., Kawaguchi, K., 2001. Growth of larval and juvenile (October 1998), 1–6.

Moku, M., Kawaguchi, K., Watanabe, H., Ohno, A., 2000. Feeding habits of three dominant myctophid fishes, Diaphus theta, Stenobrachius leucopsarus and S. nannochir, in the subarctic and transitional waters of the western North Pacific. Marine Ecology Progress Series 207, 129–140.

Munchow, A., 2000. Wind Stress Curl Forcing of the Coastal Ocean near Point Conception, California. Journal of Physical Oceanography 30 (6), 1265–1280.

Omori, M., Gluck, D., 1979. Life history and vertical migration of the pelagic shrimp Sergestes similis off the Southern California coast. Fishery Bulletin 77 (1), 183–198.

Opdal, A. F., Godo, O.-R., Bergstad, O. A., Fiksen, O., Fiksen, b. U., 2008. Distribution, identity, and possible processes sustaining meso-and bathy-pelagic scattering layers on the northern Mid-Atlantic Ridge. Deep-Sea Research II 55 (1-2), 45–58.

Pauly, D., Christensen, V., 1995. Primary production required to sustain global fisheries. Nature 374 (6519), 255–257.

Paxton, J. R., 1967. A distributional analysis for the lanternfishes (family Myctophidae) of the San Pedro Basin, California. Copeia 1967 (2), 422–440.

Pearcy, W. G., Forss, C. A., 1969. The oceanic shrimp Sergestes similis off the Oregon coast. Limnology and Oceanography 14, 755–765.

Pennington, J. T., Chavez, F. P., 2000. Seasonal fluctuations of temperature, salinity, nitrate, chlorophyll and primary production at station H3/M1 over 1989-1996 in Monterey Bay, California. Deep-Sea Research II 47 (56), 947–973.

Phillips, A. J., Brodeur, R. D., Suntsov, A. V., 2009. Micronekton community structure in the epipelagic zone of the northern California Current upwelling system. Progress in Oceanography 80 (1-2), 74–92.

Pilskaln, C. H., Lehmann, C., Paduan, J. B., Silver, M. W., 1998. Spatial and temporal dynamics in marine aggregate abundance, sinking rate and flux: Monterey Bay, central California. Deep-Sea Research II 45 (8-9), 1803–1837.

Platt, T., Denman, K. L., 1975. Spectral analysis in ecology. Annual Review of Ecology and Systematics 6 (1975), 189–210.

R Development Core Team, 2014. R: A language and environment for statistical computing.

Rebstock, G. A., 2003. Long-term change and stability in the California Current System: lessons from CalCOFI and other long-term data sets. Deep-Sea Research II 50 (14-16), 2583–2594.

Robison, B. H., 2004. Deep pelagic biology. Journal of Experimental Marine Biology and Ecology 300, 253–272.

Robison, B. H., Bailey, T. G., 1981. Sinking rates and dissolution of midwater fish fecal matter. Marine Biology 65 (2), 135–142.

Robison, B. H., Reisenbichler, K. R., Sherlock, R. E., 2005. Giant Lar-vacean Houses: Rapid Carbon Transport to the Deep Sea Floor. Science 308 (5728), 1609–1611.

Robison, B. H., Reisenbichler, K. R., Sherlock, R. E., Silguero, J. M. B., Chavez, F. P., 1998. Seasonal abundance of the siphonophore, Nanomia bijuga, in Monterey Bay. Deep-Sea Research II 45 (8-9), 1741–1751.

Robison, B. H., Sherlock, R. E., Reisenbichler, K. R., 2010. The bathypelagic community of Monterey Canyon. Deep-Sea Research II 57 (16), 1551–1556.

Roesler, C. S., Chelton, D. B., 1987. Zooplankton variability in the California Current, 1951-1982. CalCOFI Reports 28, 59–96.

Rosenfeld, L. K., Schwing, F. B., Garfield, N., Tracy, D. E., 1994. Bifurcated flow from an upwelling center: a cold water source for Monterey Bay. Continental Shelf Research 14 (9), 931–964.

Røstad, A., Kaartvedt, S., 2013. Seasonal and diel patterns in sedimentary flux of krill fecal pellets recorded by an echo sounder. Limnology and Oceanography 58 (6), 1985–1997.

Sassa, C., Kawaguchi, K., Kinoshita, T., Watanabe, C., 2002. Assemblages of vertical migratory mesopelagic fish in the transitional region of the western North Pacific. Fisheries Oceanography 11 (4), 193–204.

Schneider, D. C., Piatt, J. F., 1986. Scale-dependent correlation of seabirds with schooling fish in a coastal ecosystem. Marine Ecology Progress Series 32, 237–246.

Schwing, F. B., O’Farrell, M., Steger, J. M., Baltz, K., 1996. Coastal upwelling indices: West Coast of North America 1946-95. NOAA Technichal Memorandum NOAA-TM-NMFS-SWFSC-231, U.S. Department of Commerce.

Service, S. K., Rice, J. A., Chavez, F. P., 1998. Relationship between physical and biological variables during the upwelling period in Monterey Bay, CA. Deep-Sea Research II 45 (8-9), 1669–1685.

Skogsberg, T., Phelps, A., Society, A. P., 1946. Hydrography of Monterey Bay, California. Thermal Conditions, Part II (1934-1937). Proceedings of the American Philosophical Society 90 (5), 350–386.

Smith, S. E., Adams, P. B., 1988. Daytime surface swarms of Thysanoessa spinifera (Euphausiacea) in the Gulf of the Farallones, California. Bulletin of Marine Science 42 (1), 76–84.

Stanton, T. K., Chu, D., Wiebe, P. H., 1996. Acoustic scattering characteristics of several zooplankton groups. ICES Journal of Marine Science 53, 289–295.

Stanton, T. K., Chu, D., Wiebe, P. H., 1998. Sound scattering by several zooplankton groups. II. Scattering models. The Journal of the Acoustical Society of America 103 (1), 236–253.

Steinberg, D. K., Silver, M. W., Pilskaln, C. H., Coale, S. L., Paduan, J. B., 1994. Midwater zooplankton communities on pelagic detritus (giant lar-vacean houses) in Monterey Bay, California. Limnology and Oceanography 39 (7), 1606–1620.

Urmy, S. S., Horne, J. K., Barbee, D. H., 2012. Measuring the vertical distributional variability of pelagic fauna in Monterey Bay. ICES Journal of Marine Science 69 (2), 184–196.

Wang, D.-P., Mooers, C. N. K., 1976. Coastal-Trapped Waves in a Continuously Stratified Ocean.

Wang, Z., DiMarco, S. F., Ingle, S., Belabbassi, L., Al-Kharusi, L. H.., 2014. Seasonal and annual variability of vertically migrating scattering layers in the northern Arabian Sea. Deep-Sea Research Part I: Oceanographic Research Papers 90, 152–165.

Warren, J., 2001. In situ measurements of acoustic target strengths of gas-bearing siphonophores. ICES Journal of Marine Science 58 (4), 740–749.

Wilson, C. D., Guttormsen, M. A., 1997. Echo Integration-Trawl Survey of Pacific Whiting, Merluccius productus, off the West Coasts of the United States and Canada During July-September 1995. NOAA Technical Memorandum NMFS-AFSC-74, U.S. Department of Commerce.

Wilson, M. T., Jump, C. M., Buchheister, A., 2009. Ecology of small neritic fishes in the western Gulf of Alaska. II. Consumption of krill in relation to krill standing stock and the physical environment. Marine Ecology Progress Series 392, 239–251.

Yasuma, H., Sawada, K., Ohshima, T., Miyashita, K., Aoki, I., 2003. Target strength of mesopelagic lanternfishes (family Myctophidae) based on swimbladder morphology. ICES Journal of Marine Science: Journal du Conseil 60 (3), 584–591.

Yasuma, H., Sawada, K., Takao, Y., Miyashita, K., Aoki, I., 2010. Swim-bladder condition and target strength of myctophid fish in the temperate zone of the Northwest Pacific. ICES Journal of Marine Science: Journal du Conseil 67 (1), 135–144.

Yeh, J., Drazen, J. C., 2011. Baited-camera observations of deep-sea megafaunal scavenger ecology on the California slope. Marine Ecology Progress Series 424, 145–156.

